# Honey bee flights under ethanol-exposure show changes in body and wing kinematics

**DOI:** 10.1101/2022.01.18.476777

**Authors:** Ishriak Ahmed, Charles I. Abramson, Imraan A. Faruque

## Abstract

Flying social insects can provide model systems for inflight interactions in computationally-constrained aerial robot swarms, whose interactions may be chemically modulated under recent measurement advancements provide a capability to simultaneously make precise measurements of insect wing and body motions. This paper presents the first quantitative measurements of ethanol-exposed honey bee flight body and wing kinematic parameters. Four high speed cameras (9000 fps) were used to track the wing and body motions of insects (*Apis mellifera*). Digitization, consisting of data association, hull reconstruction, and segmentation, achieved the first high speed measurements of ethanol exposed honey bees’ wing and body motions. Kinematic changes induced by exposure to ethanol concentrations from 0% to 5% were studied using statistical analysis tools. Analysis considered trial wide mean and maximum values and gross wingstroke parameters, and tested deviations for statistical significance using Welch’s t-test and Cohen’s d test. The results indicate a decrease in maximal heading and pitch rates of the body, and that roll rate is affected at high concentrations (5%). The wingstroke effects include a stroke frequency decrease, stroke amplitude increase, stroke inclination angle increase, and a more planar wingstroke. These effects due to ethanol exposure are valuable tools to separate from interaction effects.

## Introduction

Individual insects flying in crowded assemblies perform complex aerial maneuvers by small changes in their wing motions. The complex behaviors and social interactions of honey bees (*Apis mellifera*) make them good candidates for quantifying the individual feedback rules that govern in-flight social interactions between animals. These mathematical rules may be a strong tool informing the design of autonomous aerial robotics swarm implementations on small, computationally-limited robotic platforms. Previous experiments have demonstrated that the degree of honey bee social interaction (and hence these in-flight interactions) may be chemically manipulated through exposure to chemicals such as isopentyl acetate, ethanol, or pheremones such as 9ODA and 9HDA [1].

This study extends prior work on chemical exposure studies in 2D terrestrial locomotion to untethered flight of honey bees by examining the in-flight wing and body kinematics effects of ethanol treatment in honey bees (which have not yet previously been quantified), and by performing statistical analyses on these kinematics relative to unexposed agents. Nineteen motion variables are tracked for each case: 15 body states and 4 gross wingstroke parameters. The analysis approach tests mean and maximum values (computed over each trial) for statistical significance using Welch’s t-test and Cohen’s d test.

## Previous work

Previous studies support the use of honey bees as a model for chemically-mediated social interactions. Bees engage in a wide range of simple and complex behaviors that include learning, communication. Honey bees foraging on fermenting nectar and fruit may naturally consume ethanol. While honey bees do not have a life stage dependent on alcohol (unlike fruit flies) [2], they readily self-administer high quantities and concentrations of alcohol [3] and demonstrate preferences for specific types of alcohol [4]. Bees and humans have been recorded exhibiting parallel aggression, locomotor, and learning changes following ethanol consumption [5,6]. Ethanol reduces the sting extension response threshold [6] and increases the number of stings [5]. High levels of exposure negatively impacts passive avoidance learning [7].

Locomotor activity decreases are dose-dependent [8], with small quantities inducing erratic movements [9] and high EtOH doses inducing decreases in both bee flight and walking activity [3,10]. Free flight foraging behaviors suggest the species can building ethanol tolerance [7].

Ethanol dose-dependent learning impairments have also been recorded in honey bees [8,11,12], even in learning tasks as simple as association between an odor (conditioned stimulus) and a sucrose reward (unconditioned stimulus) in proboscis extension response (PER) experiments e.g. [8].

The previous work indicates that general honey bee behaviour is changed under ethanol influence and their flight behaviour may potentially be impacted as well. However, a review of archival literature shows that digitized recordings of in-flight wing and body motions for ethanol-exposed honey bees have not previously been reported. In this study, high speed visual tracking is used to measure body and wing motion states in flight after consumption of concentrations from 0% to 5%, and statistical tests (Welch’s t-test, Cohen’s d effect size) are applied to those measurements to reveal these effects.

## Methods and approach

### Experimental procedure

#### Chemically-exposed honey bee preparation

Foragers exiting a research hive were captured and anesthetized via storage below 0° C for 3 minutes and restrained in a harness made from a modified micro centrifuge tube. The insects were fed sucrose solution until no PER was present and let rest for approximately 24 hours at 22° C. This preparation ensured a consistent metabolic state at the beginning of experiments [13] and minimized STRANGE effects [14]. A subset of insects were then fed sucrose-based solutions with varying ethanol concentration [8], kept for 15 minutes, and added to the flight test chamber. Each insect was removed from the test chamber less than 40 minutes after introduction to ensure flight is recorded under chemical influence.

#### High speed kinematics measurement

A transparent T-shaped tunnel was attached to an *Apis mellifera* hive entrance with the two remaining exits exiting to outdoor space. Four Photron high speed cameras filmed the T-joint intersection at 9000 Hz. The intersection was isolated with partitions in order to work as a confined 2337.80 in^3^ test volume, with an 875.67 in^3^ simultaneous capture volume. Recording was initiated manually when the insects started flying in the visible volume and ended when they left the volume covered by 3 or more cameras.

#### Digitizing tool

Recorded insect flight trajectories were digitized using a high speed visual insect swarm tracker (Hi-VISTA) [15] implemented in MATLAB which can provide high-resolution tracking of multiple insects using a multiple camera system. The Hi-VISTA tracker takes synchronized frames from different cameras, identifies and removes background to recover multiple insect “blobs” which are then associated in different views. These insect targets reconstructed through voxel carving by checking consistency in views with the aid of camera projection matrices. Using the reconstructed insect visual hull, Hi-VISTA then segments the insects into wing and body and applies principal component analysis to vector geometry to determine their poses.

### Analysis

#### Body parameters considered

For this study, the state variables in each flight sequence are represented by 15 scalar variables. For a time history over [0, *T_r_*], where *T_r_* is the time length recorded, time *t* was discretized as *t_i_*, *i* =1, 2, 3..., *n* at a constant sample frequency, and the mean value of a variable *h*(*t*) measured the flight sequence was calculated as

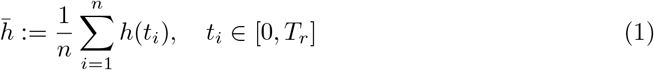

and the maximum value is defined as

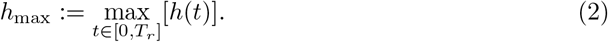

Body parameters in each flight sequence was characterized by 13 scalar values as shown in Table. 1. We define the set of these scalars as *B.* Each of these variables are measured from the stability axes of the insect. For each *s* ∈ *B* we consider population-wise mean and standard deviation.

**Table 1.** Characterizing body *B* and wing *W* variables in flight sequence.

| Set | Notation | Description |
| --- | --- | --- |
| $B$ | $\theta_b$ | Mean body pitch angle |
| | $\vert \bar{u}\vert $ | Mean absolute forward speed |
| | $\vert \bar{v}\vert $ | Mean absolute sideways speed |
| | $\vert \bar{w}\vert $ | Mean absolute heave speed |
| | $\vert \bar{p}\vert $ | Mean absolute roll rate |
| | $\vert \bar{q}\vert $ | Mean absolute pitch rate |
| | $\vert \bar{r}\vert $ | Mean absolute yaw rate |
| | $\vert u\vert _{\max}$ | Maximum absolute forward speed |
| | $\vert v\vert _{\max}$ | Maximum absolute sideways speed |
| | $\vert w\vert _{\max}$ | Maximum absolute heave speed |
| | $\vert p\vert _{\max}$ | Maximum absolute roll rate |
| | $\vert q\vert _{\max}$ | Maximum absolute pitch rate |
| | $\vert r\vert _{\max}$ | Maximum absolute yaw rate |
| $W$ | $f$ | Peak wingbeat frequency |
| | $\Phi$ | Peak wingbeat amplitude |
| | $\beta$ | Stroke plane angle |
| | $\delta$ | Dorsal bias |

The population mean value of a variable was defined as

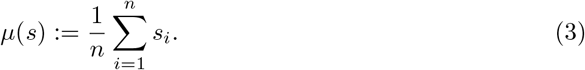

where *n* is the number of flight sequences recorded in the respective category (0%,1%,2.5%,5%,). The population standard deviation of a variable was defined as

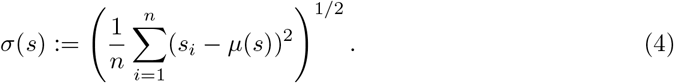

#### Wing parameters considered

The insect wingstrokes were analyzed as a set *W* comprised of 4 scalar variables.

The wing stroke, elevation, and pitch angles are represented as 3-1-2 Euler angles (*ϕ, ψ, α*). Gross stroke frequency was determined by peak to peak time difference *T_p_* = 1/*f* in *ϕ*(*t*) of left wing. *ϕ, ψ* timehistories are then resampled to have a fixed number of discrete data points *N* over each wingstroke. For each wingstroke the stroke plane angle *β* and dorsal bias *δ* is determined by fitting

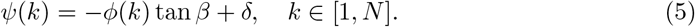

*β* can be used to compute planar motion of wing as in [16,17]

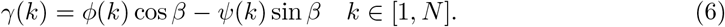

The stroke amplitude **Φ** for the wingstroke can be determined from the peak frequency of the Fourier transform of *γ*. Both wing motions were considered while determining *β, δ*, **Φ** by concatenating the datapoints.

The population mean and standard deviation of these variables *s* ∈ *W* are determined as

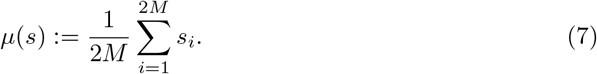

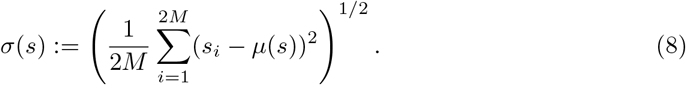

where *M* is the number of total wingstrokes recorded in the respective category (0%, 1%, 2.5%, 5%).

#### Statistical analysis tools

In order to identify variables where data showed statistical differences, binary statistical analysis was applied by dividing the data in groups

(*G*_1_: 0%, *G*_2_: 1%, *G*_3_: 2.5%, *G*_4_: 5%, *G*_5_: 1, 2.5, 5%).

The statistical tools applied to this dataset were Welch’s t-test and Cohen’s d test. Welch’s t-test tests the null hypothesis that two populations have equal means for some variable. This hypothesis was tested for each *s* ∈ *S* and *s* ∈ *W* where the null hypothesis is

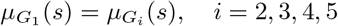

Welch’s t-test does not assume equal variance and is helpful when sample sizes are not equal. *p*-values are used to indicate the probability of the null-hypothesis being true. Cohen’s d quantifies effect size by *μ*(*s*) deviation in terms of pooled standard deviation.

## Results and discussion

In this experiment we recorded flights in bees exposed with 20% sucrose and four ethyl alcohol concentrations (by volume): 0% (Control), 1%, and 2.5% and 5%. 33 flight sequences were recorded with 33 insects (9 in 0%, 8 in 1%, 8 in 2.5%, 8 in 5% respectively). Overall, the 0%, 1%, 2.5%, and 5% trials contain 1499, 1367, 888, and 1036 wing beats, respectively.

The overall data is summarized in Table. 2, and the raw data characteristics of the affected variables are presented in Fig. 1 and 2. The following sections detail ethanol-related reductions in maximum heading and pitch rates, decrease in wing frequency and loop size, and increase in amplitude and inclination angle.

**Table 2.**
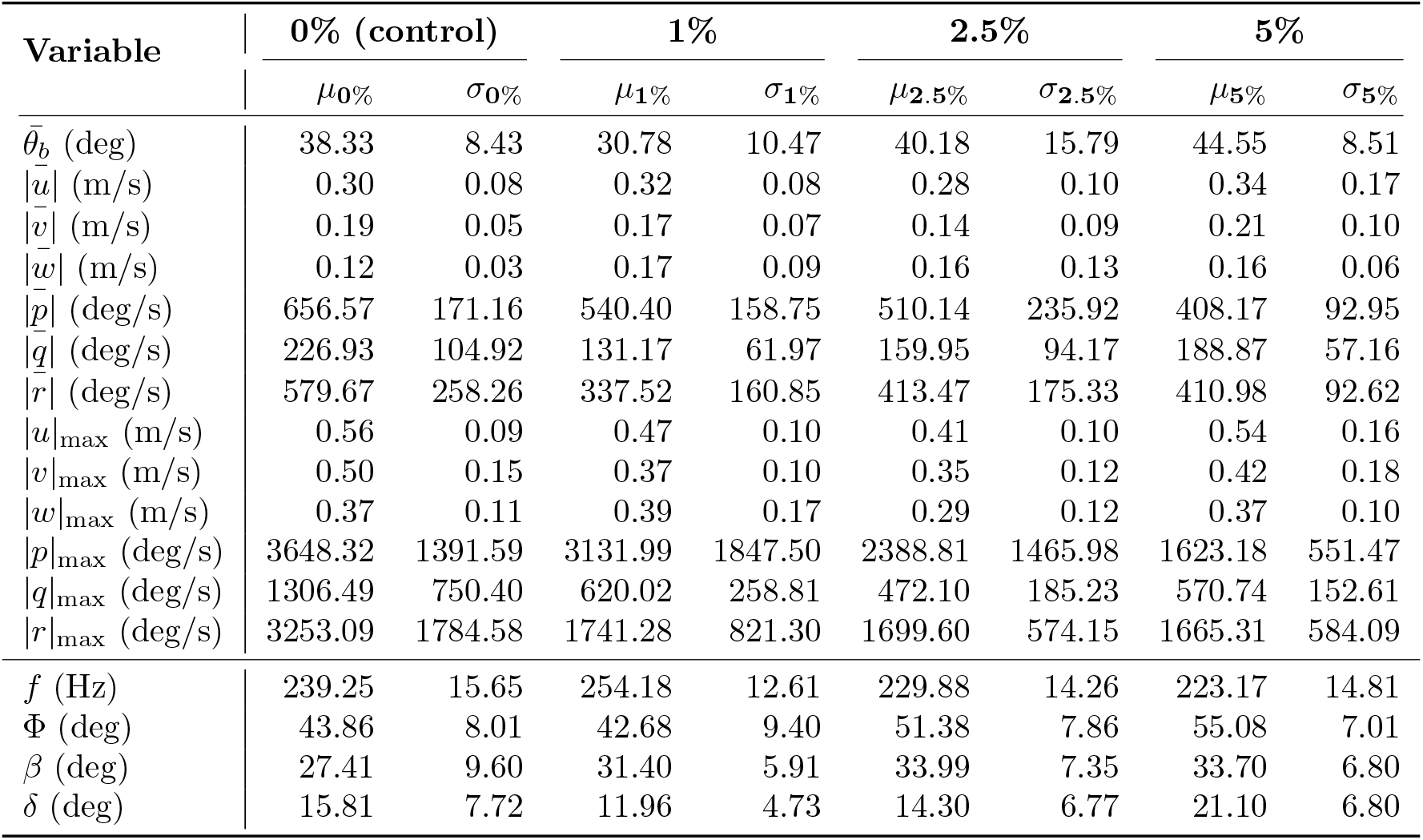
Mean *μ_i_* and standard deviation *σ_i_* of *i* =[0%, 1%, 2.5%, 5%] concentration datasets.

**Fig 1.**
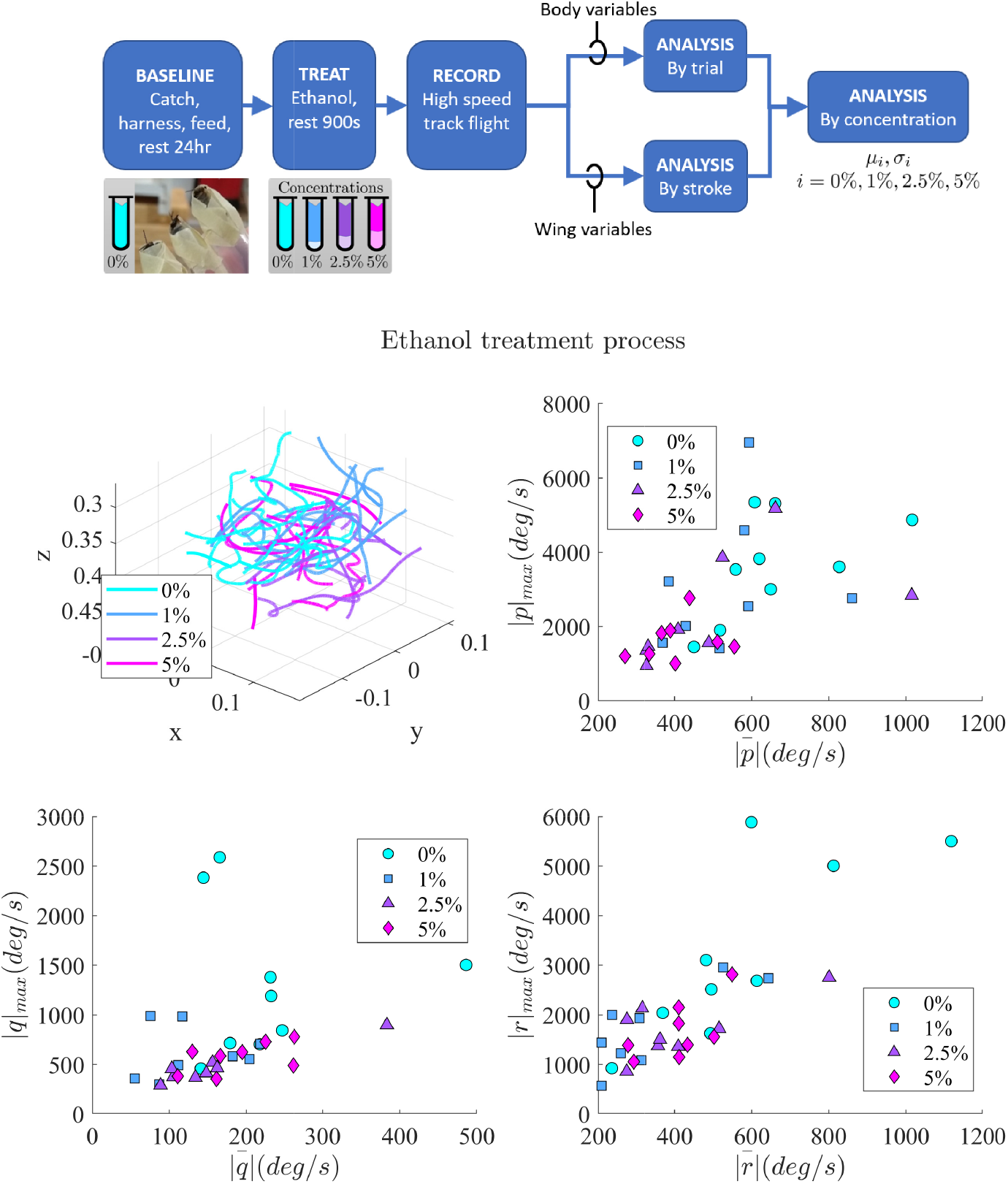
Experimental procedure and body variable characteristics. A flowchart shows the experimental process. 3D flight trajectories show the untethered flights analyzed. Maximum vs. average body rotation rates for roll (*p*), pitch (*q*), and heading (*r*). Maximal heading and pitch rate each decrease with ethanol exposure. Roll rate is affected in 5% concentration.

**Fig 2.**
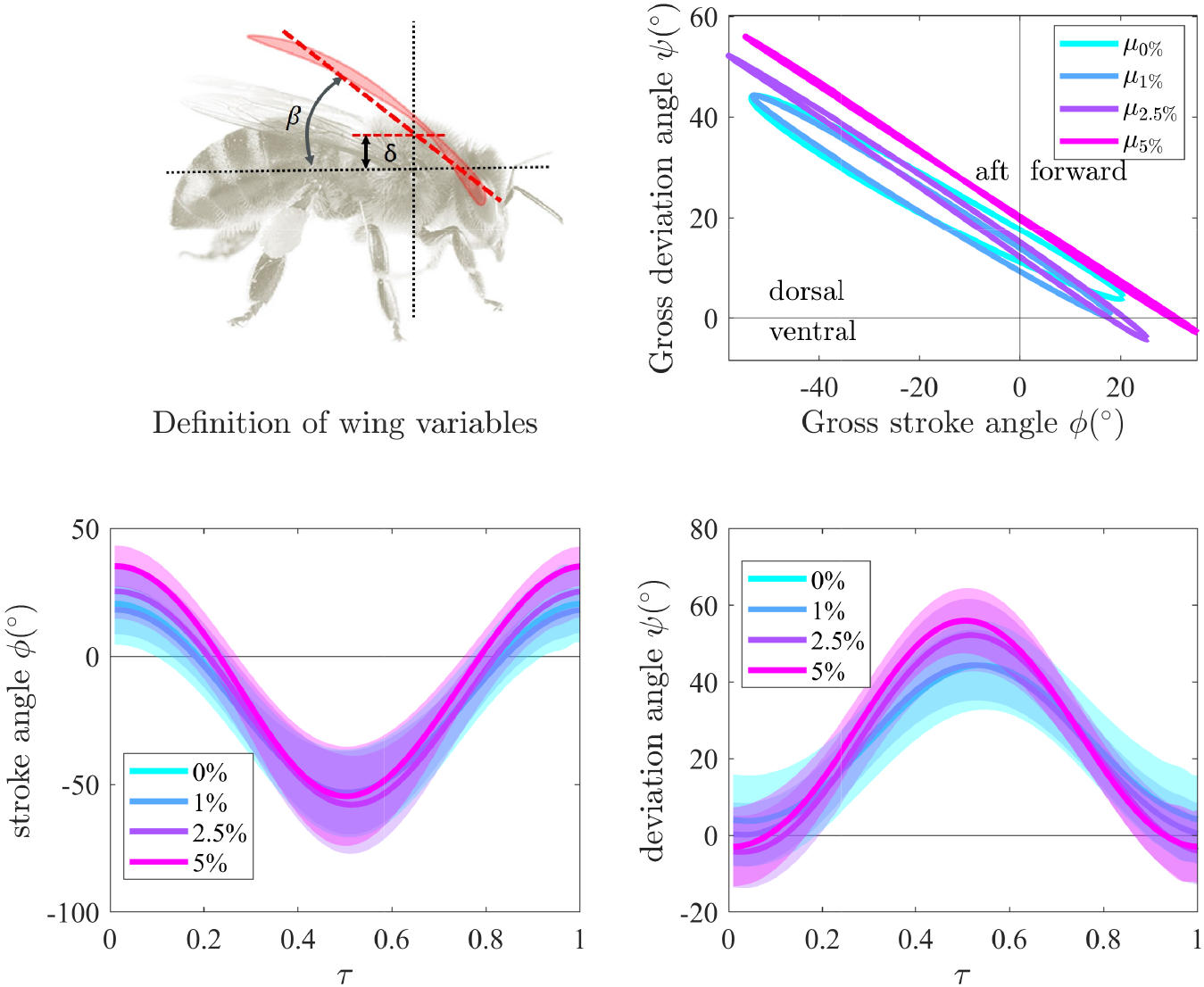
Affected wing variables. Stroke vs deviation angle of each concentration’s mean wingstroke pattern (top) shows the increasingly planar strokes with concentration. Wing stroke and deviation angle time histories (bottom 2 plots) are shown as a mean stroke *μ* over all wingbeats at that concentration ± 1 standard deviation *σ* for each concentration indicate the changes exceed standard deviation.

### Body variable characteristics

#### Pitch rates: 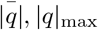

Mean absolute pitch rates decreased in all percentages but the decrease is significant only in the 1% case (*p* = 0.03, *d* = −1.09). Maximum absolute pitch rate |*q*|_max_ is significantly reduced in all comparisons (*G*_2_, *G*_3_, *G*_4_, *G*_5_) with (*d* < −0.8). Overall, The changes in maximum pitch rates between control and exposed group (G_5_) have a large Cohen-d effect size with *p* = 0.01, *d* = −1.79. A number of previous analyses indicate that airframe pitch modes are often unstable without neural feedback [18–20], this shift could signify that the pitch rate control mechanisms may have been affected and the unexposed insects flight envelopes include more aggressive motions.

#### Heading rate 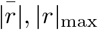

The overall decrease in mean absolute heading rates is significant only in the 1% case (*p* = 0.03, *d* = −1.09). Maximum absolute heading rate |*r*|_max_ is significantly reduced in all comparisons (*G*_2_, *G*_3_, *G*_4_, *G*_5_), with a large Cohen-d effect sizes (*d* < −0.8). Previous work has indicated that “flapping counter-torque” provides passive stabilization through aerodynamic damping on this axis [21,22], suggesting the reduction in maximum heading rate may have different interactions with the underlying airframe relative to pitch rate.

#### Roll rate: 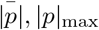

Both the maximum and mean absolute roll rates show that they are significantly reduced in the 5% case compared to control bees and not significantly in 1% and 2.5% groups. However, the overall comparison of control and exposed group (*G*_5_) show reduction in both 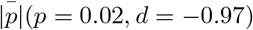, |*p*|_max_(*p* = 0.03, *d* = −0.87).

#### Body Speeds

The mean body speeds are unaffected over the dataset in every comparison. 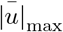 and 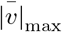 had reduced in 2.5% to significantly affect the control vs exposed case. This trend is not continued in 5% case and cannot be conclusively linked to ethanol exposure.

### Gross wingstroke characteristics

Insect asynchronous flight muscles generally operate near mechanical resonance [23] and frequency deviations often result in reduced performance. Hovering honey bees have primarily use relatively short amplitude and high frequency wingstrokes and maneuver via amplitude tuning [24,25]. Compared to control honey bees we see frequency increase (*p* << 0.05, *d* = 1.04) and amplitude decrease (*p* << 0.05, *d* = −0.13) in 1% group. On the contary, frequency decrease (*p* << 0.05, *d* = −0.61 and *d* = −1.04) with amplitude increase (*p* << 0.05, *d* = 0.94 and *d* = 1.47) in 2.5% and 5% groups. The 1% bees does not follow the trend of decreasing frequency with increased exposure level. Frequency decrease and amplitude increase was also observed in a separate dataset with manual determination of frequency by video footage observation [13].

Stroke plane inclination *β* increased in every ethanol-exposed groups (1%, 2.5%, 5%) with a medium effect size (*p* << 0.05, *d* = 0.50,0.74,0.73). *β* as a control input affects both forward flight speed *u* and pitch rate *q* [26]. Although increasing *β* tends to increase forward speed, this effect can be mitigated by changes in flight force due to the frequency and amplitude changes. The dorsal bias *δ* decreased significantly in the 1% group (*p* << 0.05, *d* = −0.59) but it increased significantly (*p* << 0.05, *d* = 0.71) in 5% group and thus shows no clear trend. A gradual mean loop size decrease with ethanol concentration (i.e., more planar wingstroke) is also visible in Fig. 2 (top). The mechanics and effect of non-planar wingstrokes are still not well understood [17,27,28] and require further aerodynamic analysis.

### Limitations and observations

Honey bees exposed to 10% ethanol solution did not initiate flight within 60 minutes in the test chamber and this study did not consider their flights. The 2.5% and 5% subjects initiated flight within 30 minutes of introduction to the test volume and displayed erratic ground movements prior to flight. Trials conducted at the 1% concentration in free-flight are quantitatively different from control insects and are distinct from the effects at higher concentrations, suggesting ethanol treatment effects are not a simple monotonic trend.

Statistical analyses such as these are limited to quantifying effects and the relative likelihood of such a measurement occurring due to chance. They do not identify the physiological or neural mechanisms behind such effects. This analysis also does not account for inter-dependence of variables. These are the first recorded quantitative high speed measurements of ethanol exposed honey bee flight, and experimental limitations on the number of animals constrain the dataset size. The differences were indicated as statistically significant and persisted when individual wingstrokes were analyzed (versus trial wide analysis), a combination which strengthens the study’s applicability.

## Conclusion

This paper presents the first quantitative high speed measurements of ethanol-exposed honey bee flight body and wing kinematic parameters. Kinematic changes induced by exposure to ethanol concentrations from 0% to 5% were studied using statistical analysis tools. The maximum heading and pitch rates reduce with increased ethanol exposure, while roll rates were affected at the 5% exposure level. Wingstroke analysis indicates a frequency decrease and amplitude increase for greater than 1% percentage exposure. Wingstroke loop size decreased and wing inclination angle increased with increased exposure level. Understanding the flight variables induced by this chemical manipulation in non-interacting flight conditions is an important result to distinguishing the effects of chemically mediated social interactions with neighboring flyers from chemical effects themselves. The study of flight behavior of honey bees to ethanol may also help us to better understand the mechanics of fine motor activity under the influence of ethanol.

## Acknowledgments

This work was supported in part by ONR N00014-19-1-2216, NSF 1950805 and NSF 1743753.

